# scGAD: single-cell gene associating domain scores for exploratory analysis of scHi-C data

**DOI:** 10.1101/2021.10.22.465520

**Authors:** Siqi Shen, Ye Zheng, Sündüz Keleş

## Abstract

**Summary:** Quantitative tools are needed to leverage the unprecedented resolution of single-cell high-throughput chromatin conformation (scHi-C) data and to integrate it with other single-cell data modalities. We present single-cell gene associating domain (scGAD) scores as a dimension reduction and exploratory analysis tool for scHi-C data. scGAD enables summarization at the gene level while accounting for inherent gene-level genomic biases. Low-dimensional projections with scGAD capture clustering of cells based on their 3D structures. scGAD enables identifying genes with significant chromatin interactions within and between cell types. We further show that scGAD facilitates the integration of scHi-C data with other single-cell data modalities by enabling its projection onto reference low-dimensional embeddings.

**Availability:** scGAD is part of the `BandNorm` R package at https://sshen82.github.io/BandNorm/articles/scGAD-tutorial.html.

**Supplementary information:** Supplementary data are available at *Bioinformatics* online.

## 1 Introduction

Single-cell technologies that profile chromatin conformation at the single-cell level emerged as promising approaches for high-resolution 3D genome characterization (Nagano *et al.*, 2013; Stevens *et al.*, 2017; Ramani *et al.*, 2017; Li *et al.*, 2019; Lee *et al.*, 2019; Tan *et al.*, 2021). Tools for specific scHi-C data inference tasks are gradually emerging (e.g., scHiCluster (Zhou *et al.*, 2019), scHiC Topics (Kim *et al.*, 2020), Higashi (Zhang *et al.*, 2021), BandNorm and 3DVI (Zheng *et al.*, 2021) for imputation and normalization of the sparse scHi-C data to advance *de facto* downstream analysis; SnapHiC (Yu *et al.*, 2020) for chromatin loop detection; scHiCTools (Li *et al.*, 2021) for quantifying cell-cell similarities). However, computational tools for extracting salient features of scHi-C data for integration with other single-cell data modalities, such as transcriptomics and epigenomics, are lacking. Here, we generalized the concept of gene-body associating domain (GAD) for bulk Hi-C data (Zhang *et al.*, 2020) and developed the single-cell Gene Associating Domain (scGAD) scores and an accompanying R package. scGAD scores adjust for inherent genomic biases and summarize scHi-C data at the gene level. As a result, scGAD scores facilitate integrating 3D genome and other genomic data from single-cell technologies.

## 2 Materials and methods

We utilized scHi-C and scRNA-seq data of developing mouse cortex and hippocampus (Dip-C and MALBAC-DT data of Tan *et al.* (2021), respectively) and Paired-Tag data of adult mouse brain (Zhu *et al.*, 2021) in our illustrations of scGAD. We first considered four variations of scGAD scores, namely *scGAD*_*raw*_, *scGAD*_*local*_, *scGAD*_*regression*_, and *scGAD*_*global*_, to investigate whether they can accommodate inherent gene-level genomic biases (Supplementary Fig. 1 and Supplementary Note). Here, *scGAD*_*raw*_ is defined as the raw total number of interactions *R*_*ij*_ for gene *i* in cell *j* without further normalization to illustrate the intrinsic genomic biases affecting the downstream analysis. As expected, *scGAD*_*raw*_ exhibits sequencing depth and gene length biases, where long genes and deeply sequenced cells dominate the cell clustering patterns (Supplementary Figs. 2-3). *scGAD*_*local*_ leverages target gene’s upstream and downstream regions of the same length as the target gene to estimate a gene-specific background. While this aims to adjust for the local background of the target gene, the neighboring regions might exhibit extreme sparsity, leading to zero number of interactions, or might harbor gene clusters, leading to overestimation of the background (Supplementary Figs. 2B-C). *scGAD*_*regression*_ first calibrates the sequencing depths of the cells and utilizes a Generalized Additive Model (GAM) to adjust for well known genomic biases, i.e., gene length, mappability, and GC content, with non-parametric smooth functions *s*_1_, *s*_2_, *s*_3_. Specifically, we have 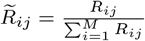,

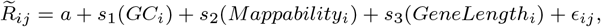

and *scGAD*_*regression*_ is the residual from this model. In contrast, *scGAD*_*global*_ removes potential gene-level biases implicitly by a standardization approach (Fig. 1A):

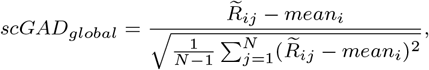

where 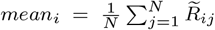. Note that gene-wise standardization of the residuals from the GAM model is operationally equivalent to standardization of 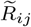, namely *scGAD*_*global*_. Hence, we kept *scGAD*_*regression*_ as unstandardized. We evaluated these variations for their performance in cell clustering and revealing relationships between the cell types. These comparisons revealed that *scGAD*_*global*_ performs the best by separating the cell types (Fig. 1B), almost as good as the full Hi-C contact matrices (Zheng *et al.*, 2021), and recovers the known cell-type relationships (Supplementary Figs. 4-5). We set *scGAD*_*global*_ as the formal definition of the scGAD scores for all the downstream analyses.

**Fig. 1.**
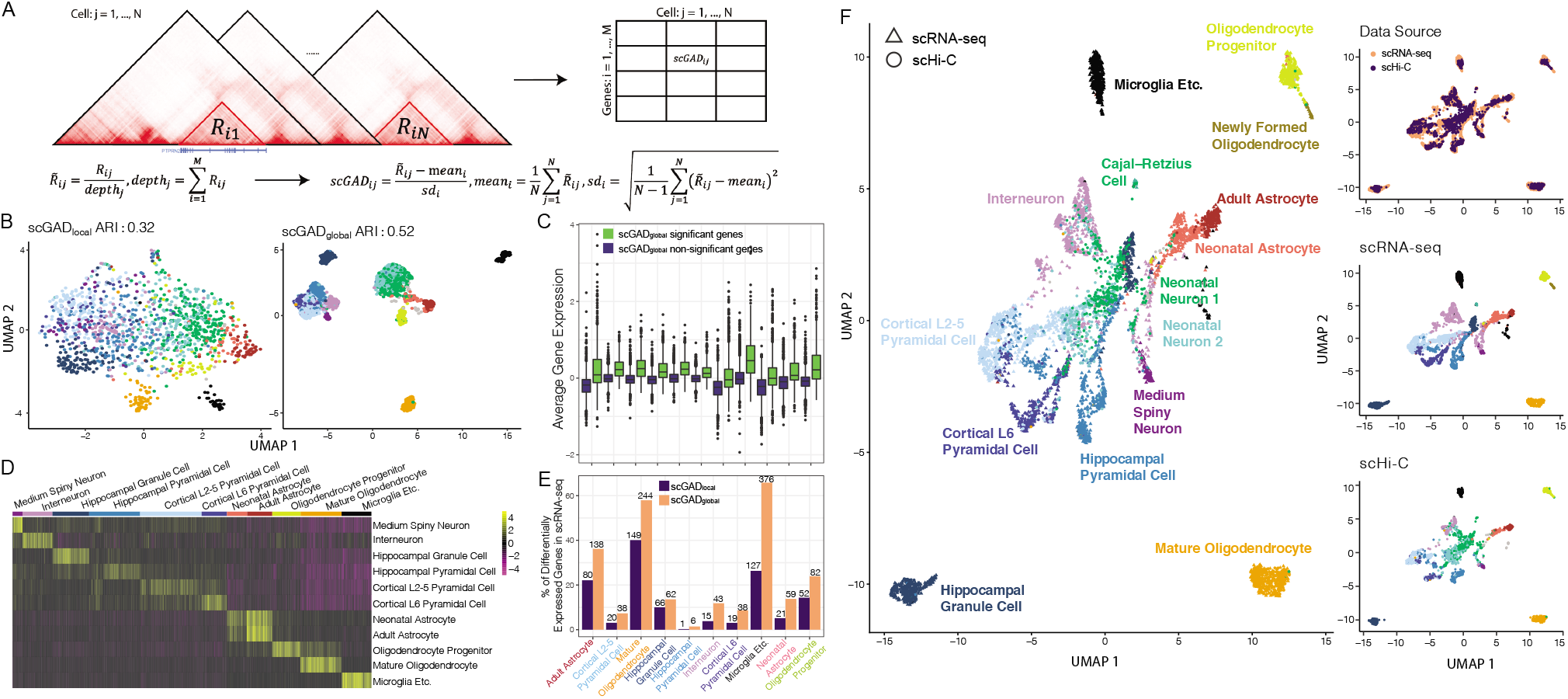
A. Overview of scGAD score calculation. *R*_*ij*_ is the total number of interactions within the promoter and body regions of gene *i* in cell *j*. This calculation corresponds to best performing *scGAD*_*global*_ and is implemented as default in the R package. B. Comparison of the low-dimensional embeddings depicting cell-type separation by different scGAD score variations. Visualizations for all the variants are available in Supplementary Fig. 4. Adjusted Rank Index (ARI) quantifies the consistency between the *k*-means clustering of the cells in the low-dimensional embeddings (with top 50 principal components) and the true cell-type labels. C. Comparison of the gene expression between significant and non-significant marker genes detected by scGAD scores across cell types shared between scHi-C and scRNA-seq data. Each gene’s scGAD score and expression are averaged over cells within each cell type. Genes with significantly high scGAD scores have significantly high gene expression (p-values from t-tests for each cell type < 10^−26^; Supplementary Fig. 7C). D. Average scGAD scores across cell-type-specific marker genes defined from scRNA-seq. Rows depict cell types in scRNA-seq, columns are the cells from the scHi-C data grouped by their cell types, and heatmap entries represent the average scores of scGAD marker genes. E. The percentage (y-axis) and the number (counts at the top of bars) of cell-type-specific marker genes defined from scRNA-seq and recovered by scGAD cell-type-specific marker gene detection. F. Projection of scHi-C data onto reference low-dimensional embeddings of scRNA-seq data with matching cell type labels.

## 3 Result

### scGAD: genes with abundant interactions

Recent studies revealed that GAD formation is a chromatin feature of highly expressed genes (Zhang *et al.*, 2020). We evaluated whether this feature is salient in the single-cell data. The correlations between the scGAD scores and expressions of genes were markedly higher, especially for the scRNA-seq marker genes, when both quantities were quantified in the same cell type compared to those in different cell types (p-value < 10^−26^, Supplementary Fig. 6). Next, we developed a permutation strategy to detect genes with significantly high scGAD scores as a means to infer highly expressed genes in a given cell type (Supplementary Note). When we evaluated genes with significantly high scGAD scores with the corresponding scRNA-seq data, they showed significantly higher expression levels compared to the genes with insignificant scGAD scores (Fig. 1C and Supplementary Fig. 7), illustrating how genes with high scGAD scores are highly expressed.

### scGAD: marker genes of cell types

We observed that cell-type-specific marker genes defined from scRNA-seq data displayed elevated scGAD scores (Supplementary Fig. 8). Furthermore, the average scGAD scores of these marker genes revealed cell-type-specific patterns (Fig. 1D and Supplementary Fig. 9). Leveraging the same marker gene detection procedure as in scRNA-seq analysis, we identified marker genes from scHi-C data with scGAD scores. We found that a large proportion of scRNA-seq marker genes overlapped with the scGAD marker genes (Fig. 1E), and the top scGAD significant marker genes yielded cell-type-specific gene expression patterns (Supplementary Fig. 10).

### scGAD: projection onto reference low-dimensional embeddings

Finally, we asked whether scGAD scores can be exploited to project cells from scHi-C data onto a given reference low-dimensional embedding (e.g., from scRNA-seq; Fig. 1F). Projection onto the scRNA-seq embedding from the same system (i.e., with the exact same cell types) revealed that the cells originating from the same cell type but quantified by different data modalities were tightly clustered. Next, taking advantage of the Paired-Tag data, which included a larger number of cell types than the scHi-C data, we observed that scGAD facilitated an accurate projection of cells onto this larger space (Supplementary Figs. 11-12). This across-modality projection enables fast cell-type annotation for 3D genomics data and promotes integrative analyses of 3D genome structure, epigenomics, and transcriptomics to decipher gene regulation mechanisms at single-cell resolution.

In summary, scGAD provides a set of analysis tools to address the pressing needs for integrating scHi-C data with other single-cell data modalities.

