## Supplementary Note and Figures for "scGAD: single-cell gene associating domain scores for exploratory analysis of scHi-C data"

#### 1 Supplementary Note

##### Data source and pre-processing

The developing mouse cortex and hippocampus single-cell Hi-C and scRNA-seq datasets (referred to as Dip-C and MALBAC-DT, respectively, in the original paper, Tan et al. 2021) were downloaded from GEO (Barrett et al. 2012) under the accession number GSE162511. The adult mouse brain cortex and hippocampus Paired-Tag data (Zhu et al. 2021) in the processed format were obtained from the web portal at `http://catlas.org/pairedTag`. GENCODE (Frankish et al. 2021) M19 was utilized to retrieve annotation data for protein-coding genes. scHi-C data were filtered following the pre-processing procedure in the original paper (Tan et al. 2021) and resulted in 1,954 cells. The contact matrices for all the intra-chromosomal interactions were binned at 10kb resolution for each cell. Genes longer than or equal to 100kb were utilized for scGAD score calculations to ensure sufficient domain area and avoid sparsity.

##### scGAD score variants

We considered four variations of scGAD scores, namely $scGAD_{raw}$, $scGAD_{local}$, $scGAD_{regression}$, and $scGAD_{global}$ with varying levels of adjustments for inherent genomic biases. A diagram summarizing their differences is provided in Supplementary Fig. 1. $scGAD_{global}$ is selected as the formal definition of scGAD score (Figure 1A).

$scGAD_{raw}$ takes as input the raw contact matrices and calculates the number of interactions within promoter and body regions of the genes. The promoter regions are defined as 1kb upstream of the transcription start sites. No further bias removal or normalization was carried out for $scGAD_{raw}$ to illustrate the inherent genomic biases that may affect the downstream analysis (Supplementary Figs. 2-3).

$scGAD_{local}$ follows the gene-body associating domain analysis of bulk-cell Hi-C data (Zhang et al. 2020), where the target gene's neighboring upstream and downstream regions of the same length as the target gene are leveraged to estimate a gene-specific background interaction signal. $scGAD_{local}$ is defined as the ratio of the total number of interactions within the gene's promoter and body region over the average number of interactions within the upstream and downstream neighboring regions. This ratio aims to remove sequencing depth and gene length biases simultaneously. However, further investigation of the $scGAD_{local}$ scores revealed two additional biases, driven mainly by the sparse nature of the scHi-C data, that severely affect the $scGAD_{local}$ calculations. First, cells with low sequencing depth are more likely to have zero interactions in the neighboring regions of the target gene (Supplementary Fig. 2B). Consequently, $scGAD_{local}$ encounters division by zero even for highly interacting gene domains. In addition, genes located at the boundary of centromeres or within highly repetitive regions of the genome may also have underestimated interactions in either upstream or downstream regions due to technical issues for dealing with repetitive regions. This, in turn, leads to underestimated background signals. Second, gene densities of the neighborhood regions also introduce bias into $scGAD_{local}$, especially for long genes (Supplementary Fig. 2C). Specifically, gene clusters in the upstream and downstream regions of the target gene lead to overestimation of the background signals and attenuate $scGAD_{local}$ scores.

$scGAD_{regression}$ first scales $scGAD_{raw}$ by the sequencing depth for each cell to account for differences in sequencing depths of the cells. The sequencing depth is defined as the sum of $scGAD_{raw}$ scores across all the eligible genes within each cell, namely $\sum_{i=1}^{M} R_{ij}$, where $i = 1, \cdots, M$ across genes and $j = 1, \cdots, N$ across cells. Subsequently, a generalized additive model (GAM) (Hastie and Tibshirani 2017) is utilized to explicitly regress out the gene length, mappability, and GC content effects. More specifically,

$$\widetilde{R}_{ij} = \frac{R_{ij}}{\sum_{i=1}^{M} R_{ij}},$$
$$\widetilde{R}_{ij} = a + s_1(GC_i) + s_2(Mappability_i) + s_3(GeneLength_i) + \epsilon_{ij},$$

where $s_1, s_2, s_3$ are smooth, non-parametric functions that quantify the non-linear relationships between scGAD scores and gene length, mappability, and GC content. $scGAD_{regression}$ scores are the residuals from this GAM fit.

$scGAD_{global}$ exploits the same sequencing depth scaling strategy as $scGAD_{regression}$, followed by a standardization approach to eliminate an overall bias estimated across all the cells and make the scores comparable across genes:

$$scGAD_{global} = \frac{\widetilde{R}_{ij} - mean_i}{\sqrt{\frac{1}{N-1}\sum_{j=1}^{N}(\widetilde{R}_{ij} - mean_i)^2}},$$
$$\text{where } mean_i = \frac{1}{N}\sum_{j=1}^{N} \widetilde{R}_{ij}.$$

While the "local" strategy aims to estimate a background signal for each gene by using the gene's local neighborhood, the "global" strategy uses the mean scGAD score across all cell types to estimate a gene-specific background interaction signal. The unstandardized residuals from the GAM fit, namely $scGAD_{regression}$, does improve the cell type separation compared to $scGAD_{raw}$ and $scGAD_{local}$ (Supplementary Fig. 4), however, fails to exhibit cell-type-specific patterns for scRNA-seq marker genes (Supplementary Fig. 9). The standardized version (i.e., $scGAD_{global}$) outperforms the other variants regarding the cell-type separation, cell-type relationship recovery, and scRNA-seq marker gene recovery (Figure 1 and Supplementary Figs. 4-10). Notably, the standardization of the residuals from the GAM fit, i.e., $scGAD_{regression}$, is operationally equivalent to standardization of $\widetilde{R}_{ij}$, namely $scGAD_{global}$ with the notable exception that $scGAD_{global}$ does not require additional data acquisition on mappability and GC content for each gene in the relevant genome.

#### Evaluation of cell clustering and cell-type separation

UMAP visualizations of the gene by cell matrices from scGAD or scRNA-seq rely on an initial Principal Component Analysis (PCA) for dimension reduction. The first 50 principal components were utilized for UMAP projections onto 2-dimensional space with the `R` package `umap`. $k$-means algorithm from the `R` package `kmeans` was applied to the 50 PC latent embeddings for cell clustering. Adjusted Rand Index (ARI) (from the `R` package `mclust`) was employed to quantify the agreement between the $k$-means clustering labels and the true cell-type labels.

#### Permutation test for identifying genes with significantly high scGAD scores

We developed a permutation test to detect genes with highly abundant interactions within their gene-associating domains. More specifically, we generated a null distribution for the mean scGAD scores of genes by permuting the scGAD scores of genes within each cell 1,000 times. For each permutation, we recorded the average scGAD score of each gene across the $N$ cells and obtained gene-specific null distributions. We then calculated a p-value for each gene using these null distributions and controlled the false discovery rate at level 0.05 by the Benjamini-Hochberg procedure (Benjamini and Hochberg 1995). To assess the validity of the resulting significant genes, we leveraged gene expression from matching scRNA-seq data. Genes with significant scGAD scores had markedly higher average gene expression compared to the genes with non-significant scGAD scores (Supplementary Fig. 7A, B). The differences from $scGAD_{global}$ were particularly significant compared to $scGAD_{local}$ (Supplementary Fig. 7C). This further promotes $scGAD_{global}$ as yielding more accurate quantification of gene interaction activities and leading to better detection of genes with significantly high scGAD scores that are further supported by high scRNA-seq gene expression.

#### Identifying cell-type-specific marker genes from scGAD scores

We identified cell-type-specific marker genes from scGAD scores following the marker gene detection procedure of scRNA-seq data by Tan et al. 2021. Specifically, FindMarker function in the `Seurat` `R` package was utilized with `"logfc = 0.25"`, `"only.pos = TRUE"`.

##### Projecting scHi-C data summarized by scGAD scores onto reference low-dimensional embeddings

We leveraged the single-cell data integration strategy proposed in Stuart et al. 2019 to project scHi-C data onto a reference low-dimensional embedding from scRNA-seq. Specifically, we converted the scHi-C contact matrices into gene by cell matrices with entries corresponding to scGAD scores. The entries from both the scGAD and scRNA-seq gene by cell matrices were filtered to keep the common genes. Subsequently, scRNA-seq matrix was normalized by log transformation of Count Per Million (log CPM) using the "NormalizeData" function from the `Seurat` `R` package. For both scGAD and scRNA-seq matrices, variable features were selected using "SelectIntegrationFeatures" and the anchors were located using the "FindInetegrationAnchors" function. Finally, the processed scGAD and scRNA-seq data were passed into the "IntegrateData" function. All the functions from `Seurat` (Version 4) (Stuart et al. 2019; Y. Hao et al. 2021) were used with default parameters.


### 2 Supplementary Figures

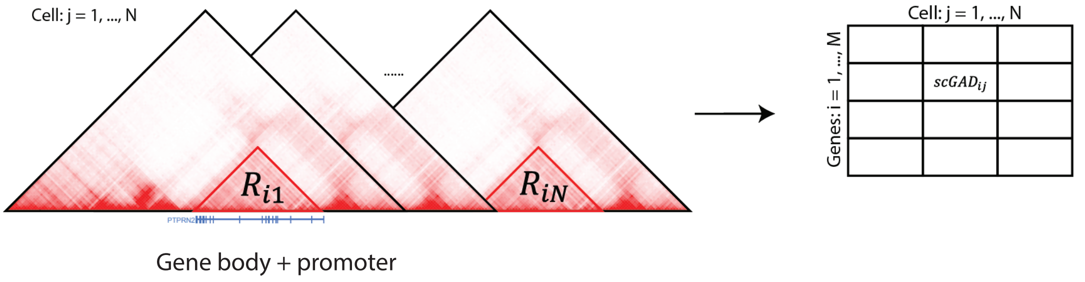


$$scGAD_{raw} = R_{ij}$$

$$scGAD_{local} =$$

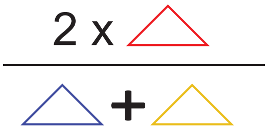


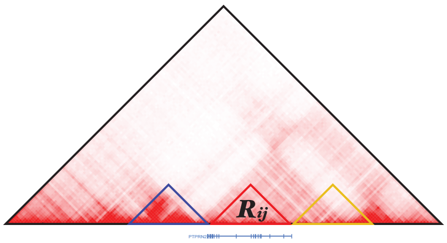


$$\widetilde{R}_{ij} = \frac{R_{ij}}{\sum_{i=1}^{M} R_{ij}}$$

$$\widetilde{R}_{ij} = a + s_1(GC_i) + s_2(Mappability_i) + s_3(GeneLength_i) + \epsilon_{ij}$$

$$scGAD_{regression} = \widetilde{R}_{ij} - \widehat{\widetilde{R}}_{ij}$$

$$mean_i = \frac{1}{N}\sum_{j=1}^{N} \widetilde{R}_{ij} \qquad sd_i = \sqrt{\frac{1}{N-1}\sum_{j=1}^{N}(\widetilde{R}_{ij} - \frac{1}{N}\sum_{j=1}^{N}\widetilde{R}_{ij})^2}$$

$$scGAD_{global} = \frac{\widetilde{R}_{ij} - mean_i}{sd_i}$$

Supplementary Fig. 1: **Single-cell Gene Associating Domain (scGAD) score variations.** $scGAD_{raw}$ for gene $i$ in cell $j$ denotes the total number of raw interactions within gene $i$'s promoter and body regions in cell $j$. $scGAD_{local}$ aims to remove the sequencing depth and gene length biases by estimating a local background interaction signal as the average interaction frequencies of upstream and downstream regions of the target gene, where the lengths of these neighboring regions are set to length of the target gene. $scGAD_{regression}$ first adjusts for sequencing depths of the cells and regresses out gene length, mappability, and GC content effects by fitting a generalized additive model (GAM) across the genes. $scGAD_{global}$ shares the same sequencing depth normalization procedure as $scGAD_{regression}$, followed by a standardization procedure to remove an overall bias estimated across all the cell types and make the scGAD scores comparable across genes and cells. $scGAD_{global}$ is utilized as the final definition of scGAD score for downstream analysis.

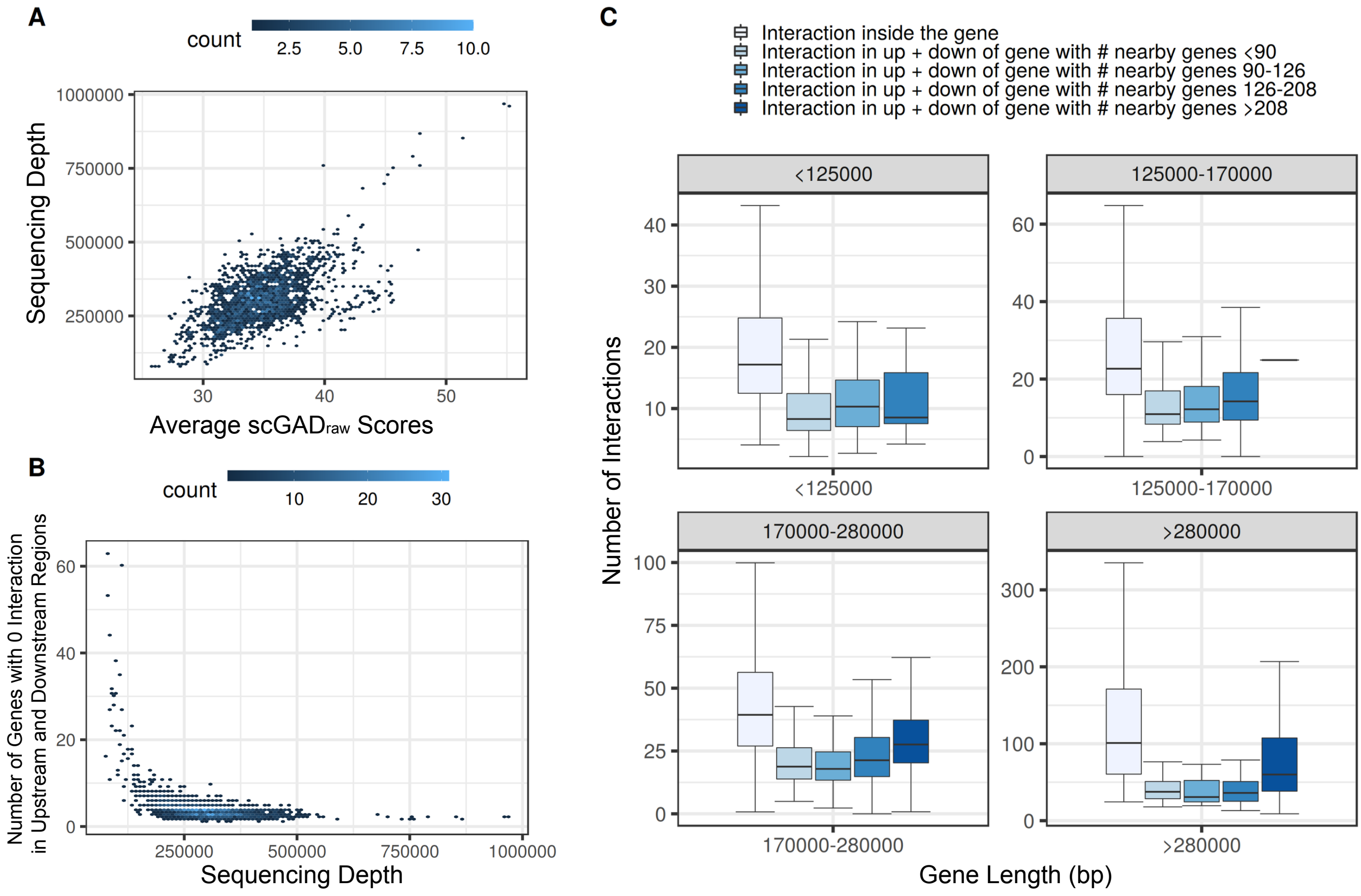


Supplementary Fig. 2: **Impact of sparsity of scHi-C data and intrinsic genomic biases on scGAD scores. A.** The average gene associating domain interaction frequencies (i.e., $scGAD_{raw}$ score without any bias correction) across genes has a strong positive correlation with the sequencing depths of the cells. x-axis denotes the average of scGAD scores across genes within a cell. Colors represent the numbers of cells in each hex bin. **B.** Number of genes with zero interaction in neighboring regions used for background estimation increases as the sequencing depth decreases. Cells with overall low sequencing depth and high sparsity tend to have a larger number of genes with 0 interaction in the upstream and downstream neighboring regions. This leads to underestimation of the background interaction signal as 0. Colors represent the numbers of cells in each hex bin. **C.** Interaction frequencies in the upstream and downstream neighboring regions with respect to the target gene lengths and the sizes of the gene clusters (gene densities of the neighboring regions). Longer genes with correspondingly long upstream and downstream neighboring regions are more likely to have larger numbers of genes in their neighborhoods. Such gene clusters tend to amplify estimated background interaction signals for $scGAD_{local}$ scores.

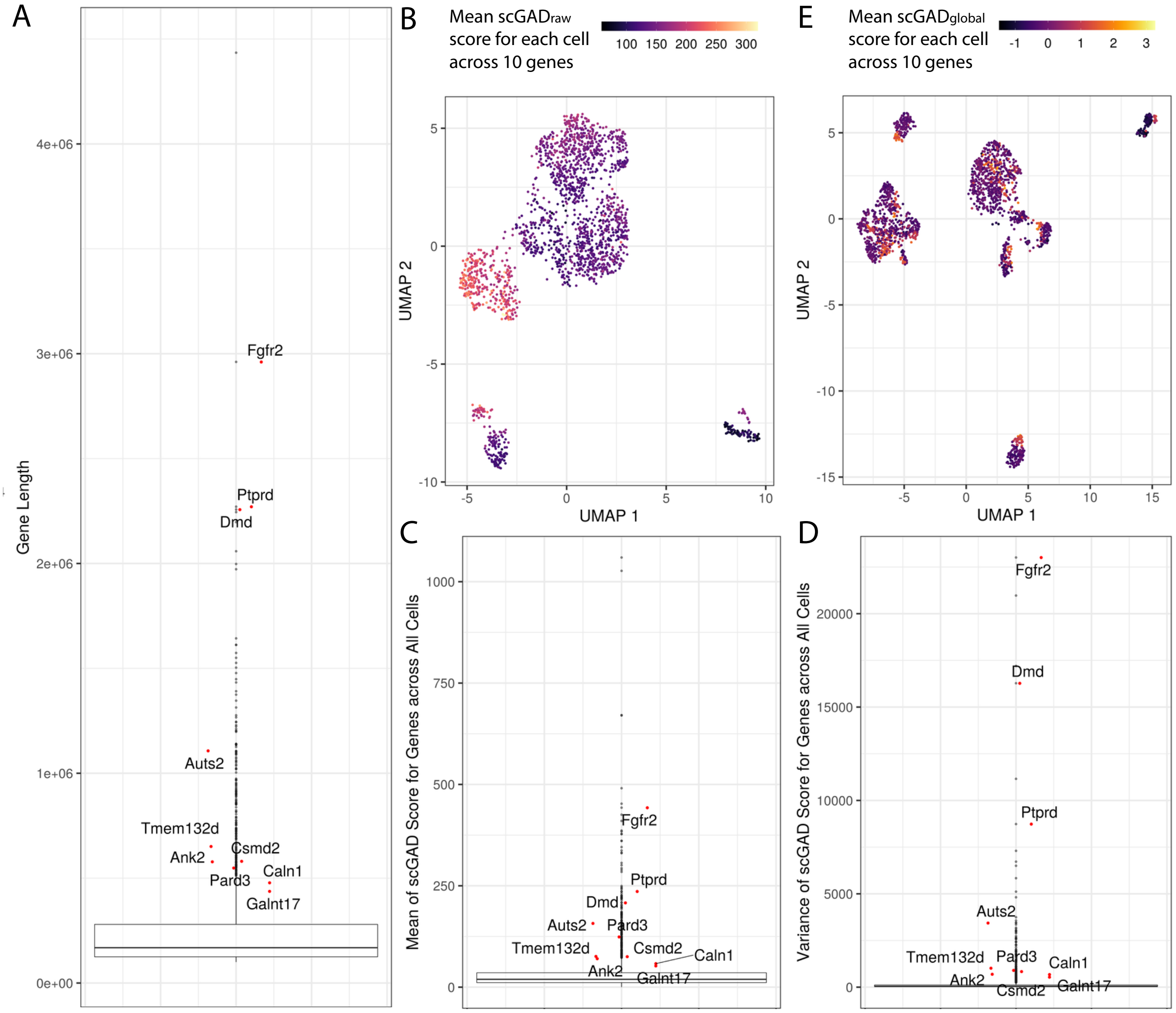


Supplementary Fig. 3: **Gene length impact on scGAD score estimation.** **A.** Principal component analysis (PCA) of $scGAD_{raw}$ revealed that the third principal component is largely correlated with the gene length. The box plot depicts the distribution of gene lengths where genes with the top ten largest (in absolute value) component loadings for the 3rd PC are marked in red. **B.** Low-dimensional projection of cells based on their $scGAD_{raw}$ with UMAP shows that the cell clustering is driven by the long genes in **A**. **C.** Average $scGAD_{raw}$ score across cells for each gene. The ten long genes in **A** exhibit markedly high average scores. **D.** Variances of $scGAD_{raw}$ scores across cells for each gene. The ten long genes in **A** exhibit markedly high variances. **E.** Low-dimensional projection of cells based on their $scGAD_{global}$ with UMAP shows that $scGAD_{global}$ standardization procedure effectively removes the gene length effect and hence the cell clustering is not driven by the long genes anymore.

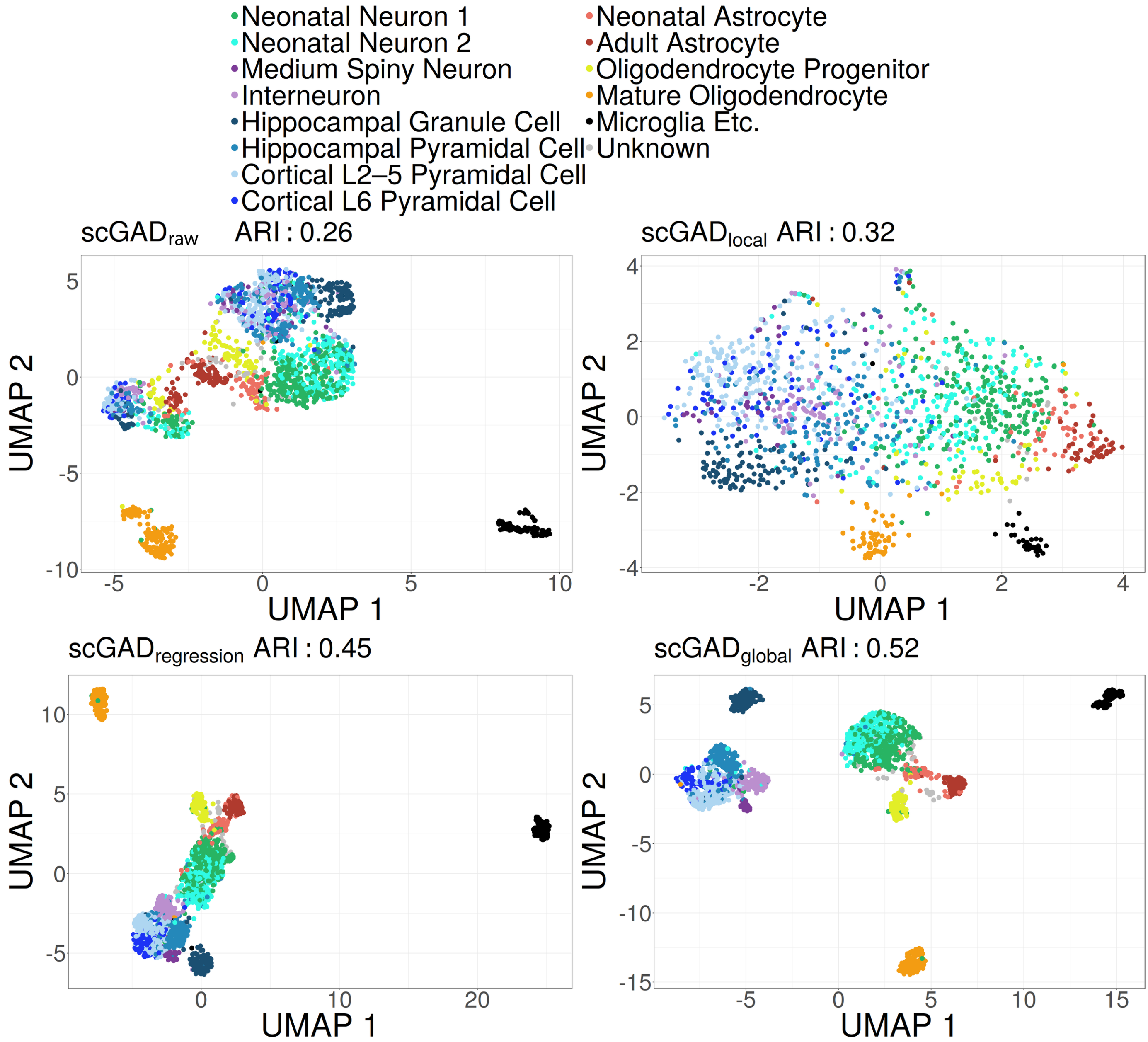


Supplementary Fig. 4: **Comparison of clustering and cell-type separation performances of scGAD score variants.** UMAP shows the low-dimensional embeddings where the cells are annotated by their true cell-type labels. Adjusted Rank Index (ARI) quantifies the consistency between the $k$-means clustering of the cells in the low-dimensional embeddings (with top 50 principal components) and the true cell-type labels.

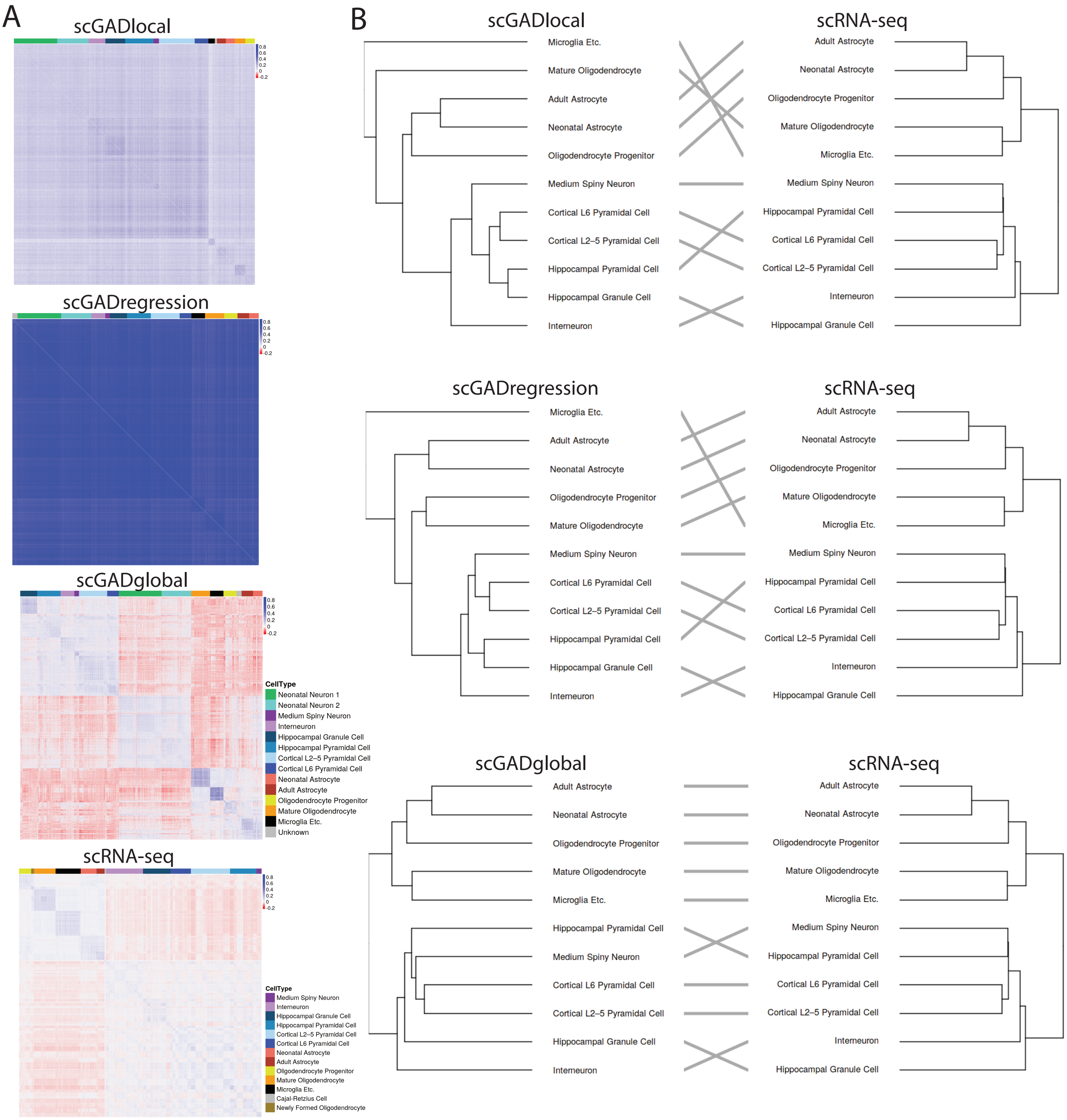


Supplementary Fig. 5: **Cell-type relationships unveiled by scGAD score variants. A.** Heatmaps of $scGAD_{local}$, $scGAD_{regression}$, $scGAD_{global}$, and scRNA-seq depict the Pearson correlation coefficients between every pair of cells with the same color scale setting. **B.** Hierarchical clustering of cell types is compared to the scRNA-seq panel, which performs as a gold standard for the cell-type relationships to be recovered by scGAD variants. The distance matrix utilized in the hierarchical cluster analysis is obtained by one minus the average of the Pearson correlation coefficients for each pair of cells between every pair of cell types. Comparison between trees of hierarchical clustering is implemented by `dendextend` R package (Galili 2015).

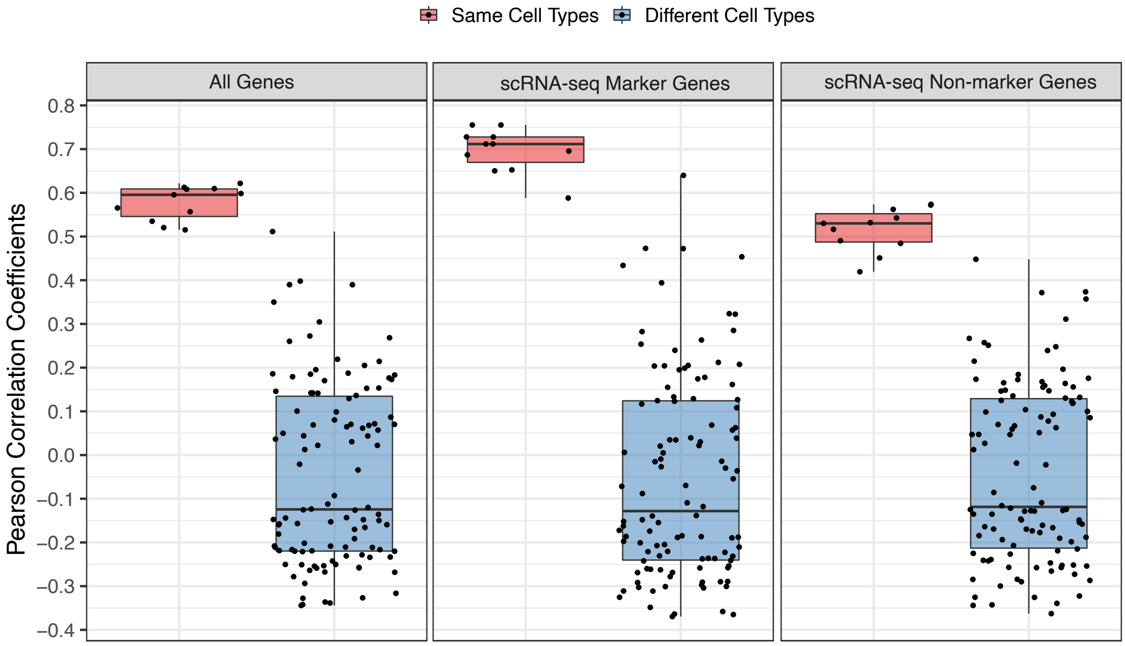


Supplementary Fig. 6: **Correlations between scRNA-seq expression and scGAD scores of genes.** For each gene, scRNA-seq expression and scGAD score are first averaged within each cell type, respectively. Then, Pearson correlation coefficients are calculated between gene scRNA-seq expression and scGAD scores within the same cell type or between two different cell types as the baseline. Genes are stratified into three categories: all the genes that are available for scRNA-seq and scGAD data, scRNA-seq marker genes, genes that are not marker genes for any cell type in the scRNA-seq data.

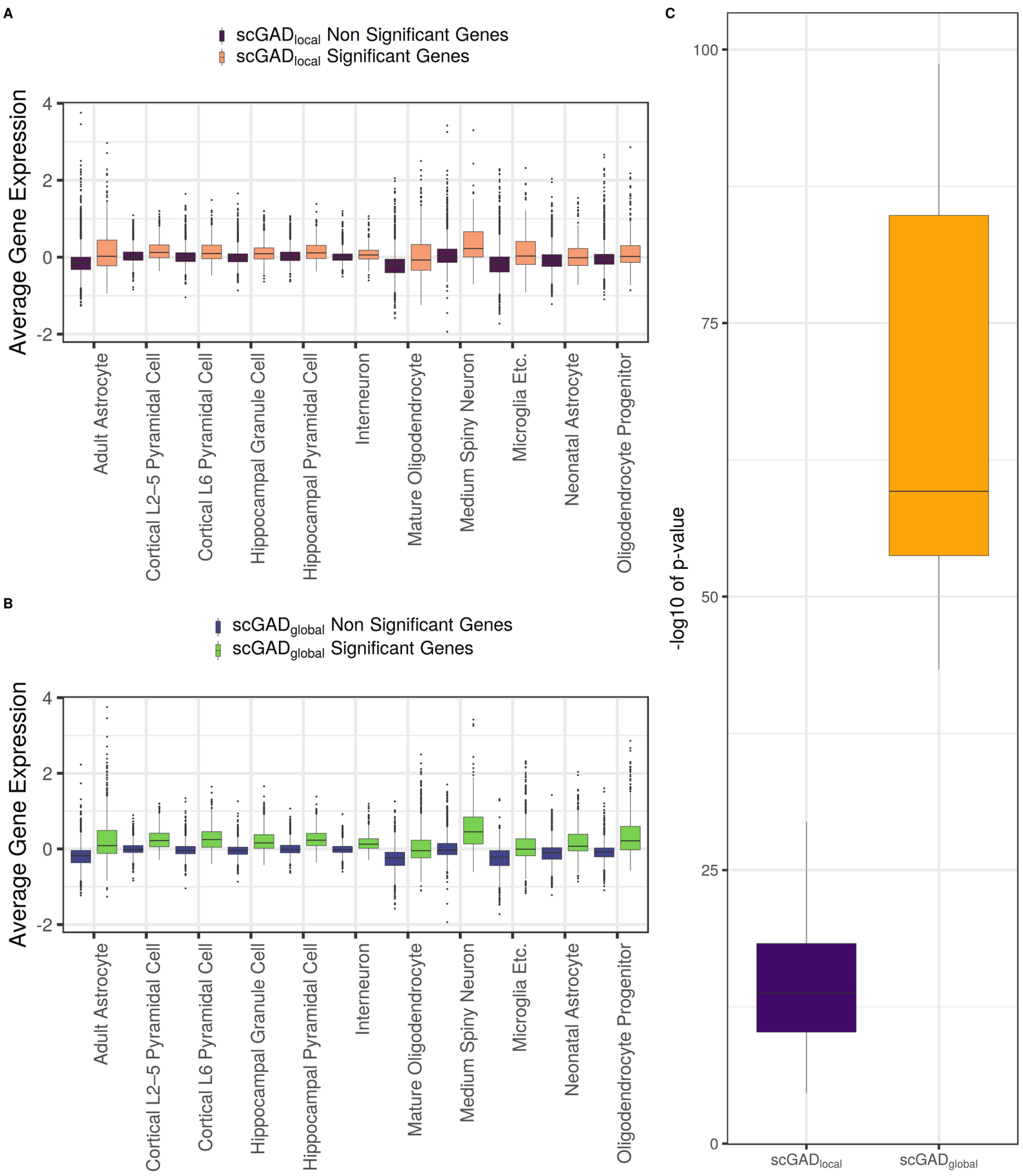


Supplementary Fig. 7: **scRNA-seq gene expression comparison between genes with and without significant scGAD scores (scGAD marker and non-marker genes). A.** Average scRNA-seq gene expression across genes that are identified as having significantly high $scGAD_{local}$ scores (orange) compared to the rest of the genes (purple) for each cell type. **B.** Average scRNA-seq gene expression across genes that are identified as having significantly high $scGAD_{global}$ scores (green) compared to the rest of the genes (blue) for each cell type. **C.** Boxplots of the $-log_{10}$ transformation of the p-values from t-tests comparing the expression of significant (scGAD marker) and insignificant (scGAD non-marker) genes for each cell type where the marker genes are defined from $scGAD_{local}$ (left boxplot) and $scGAD_{global}$ (right boxplot).

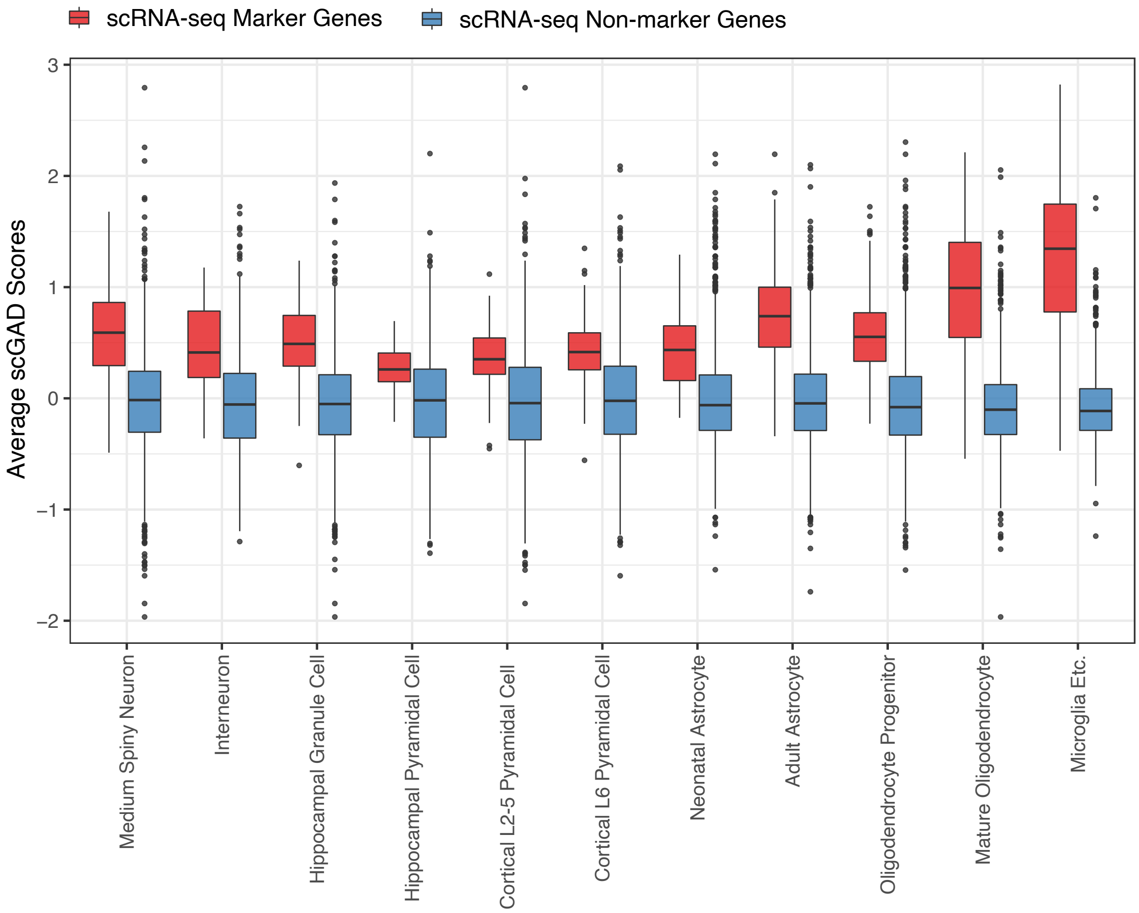


Supplementary Fig. 8: **Comparison of scGAD scores between scRNA-seq marker and non-marker genes.** Comparison of the $scGAD_{global}$ scores between cell-type-specific marker genes defined by scRNA-seq and the rest of genes for each cell type listed in the x-axis.

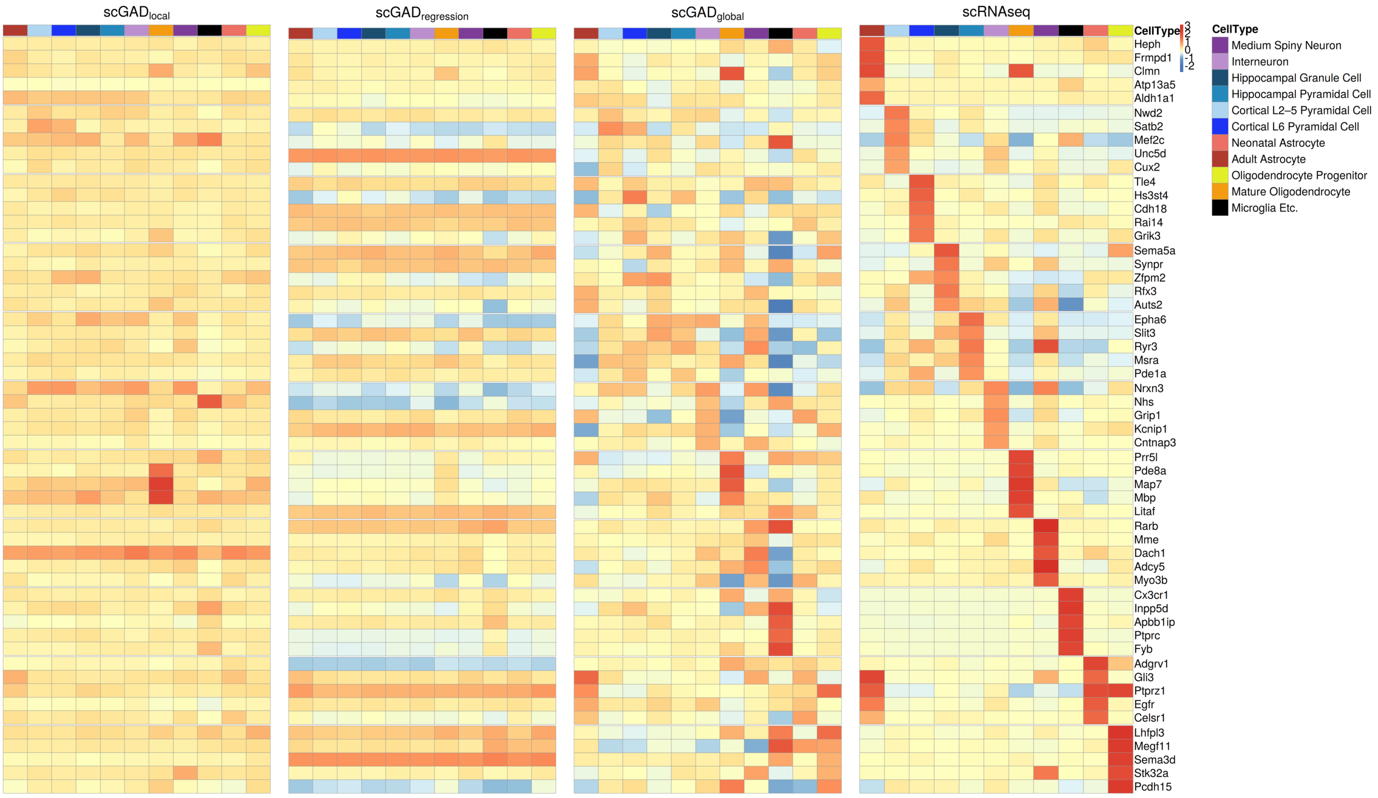


Supplementary Fig. 9: **Comparison of cell-type specific patterns between** $scGAD_{local}$**,** $scGAD_{regression}$**, and** $scGAD_{global}$ **among scRNA-seq marker gene.** Top five differentially expressed genes detected for each cell-type in scRNA-seq data (fourth panel) are utilized to visualize the cell-type-specific patterns by $scGAD_{local}$, $scGAD_{regression}$, and $scGAD_{global}$ scores. scGAD scores or gene expression are averaged across all the cells within each cell-type annotated by the color bar at the top of the heatmap.

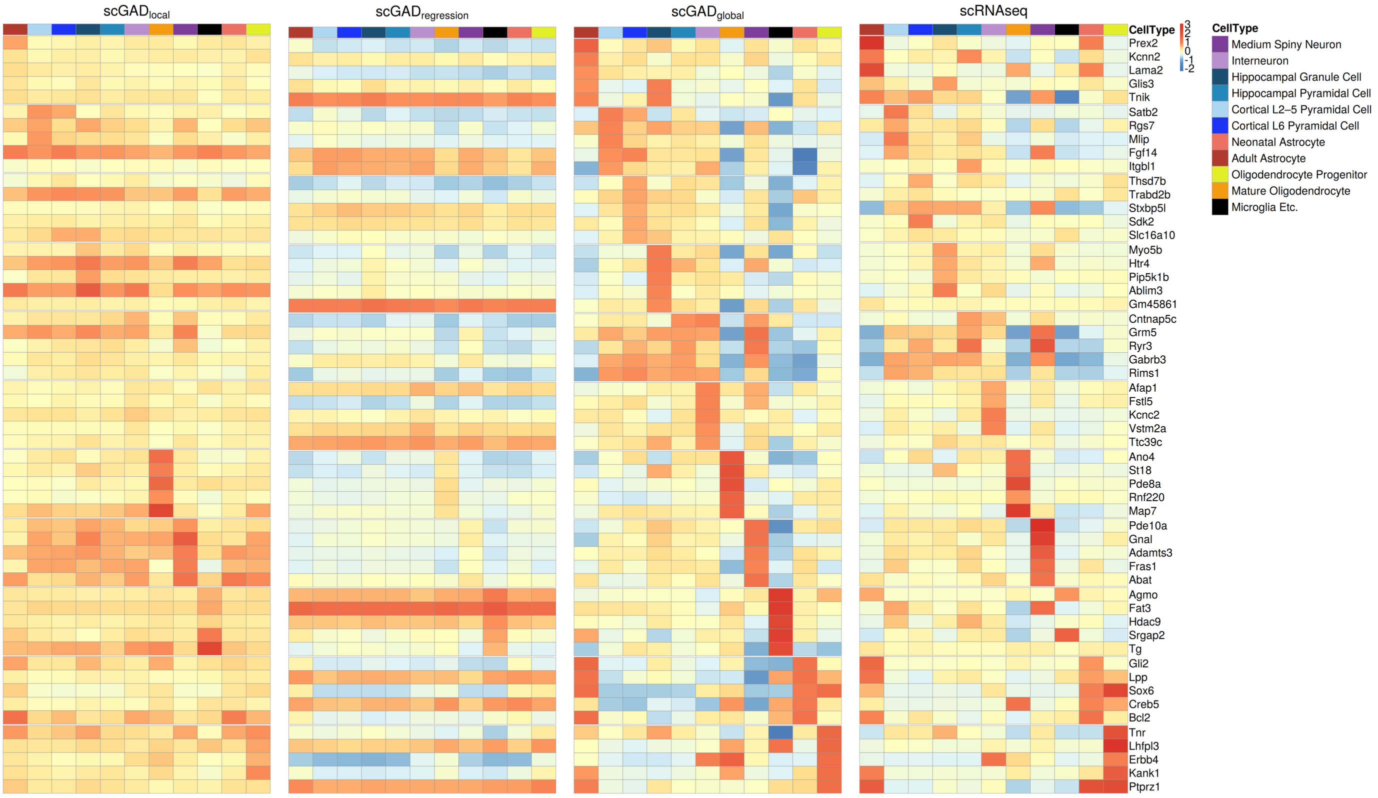


Supplementary Fig. 10: **scRNA-seq gene expression exhibits cell-type-specific patterns for scGAD marker genes.** Average cell-type-specific scRNA-seq expression of top five genes with differential $scGAD_{global}$ scores for each cell type are displayed. scGAD scores or the gene expression are averaged across all the cells within each cell-type annotated by the color bar at the top of the heatmap.

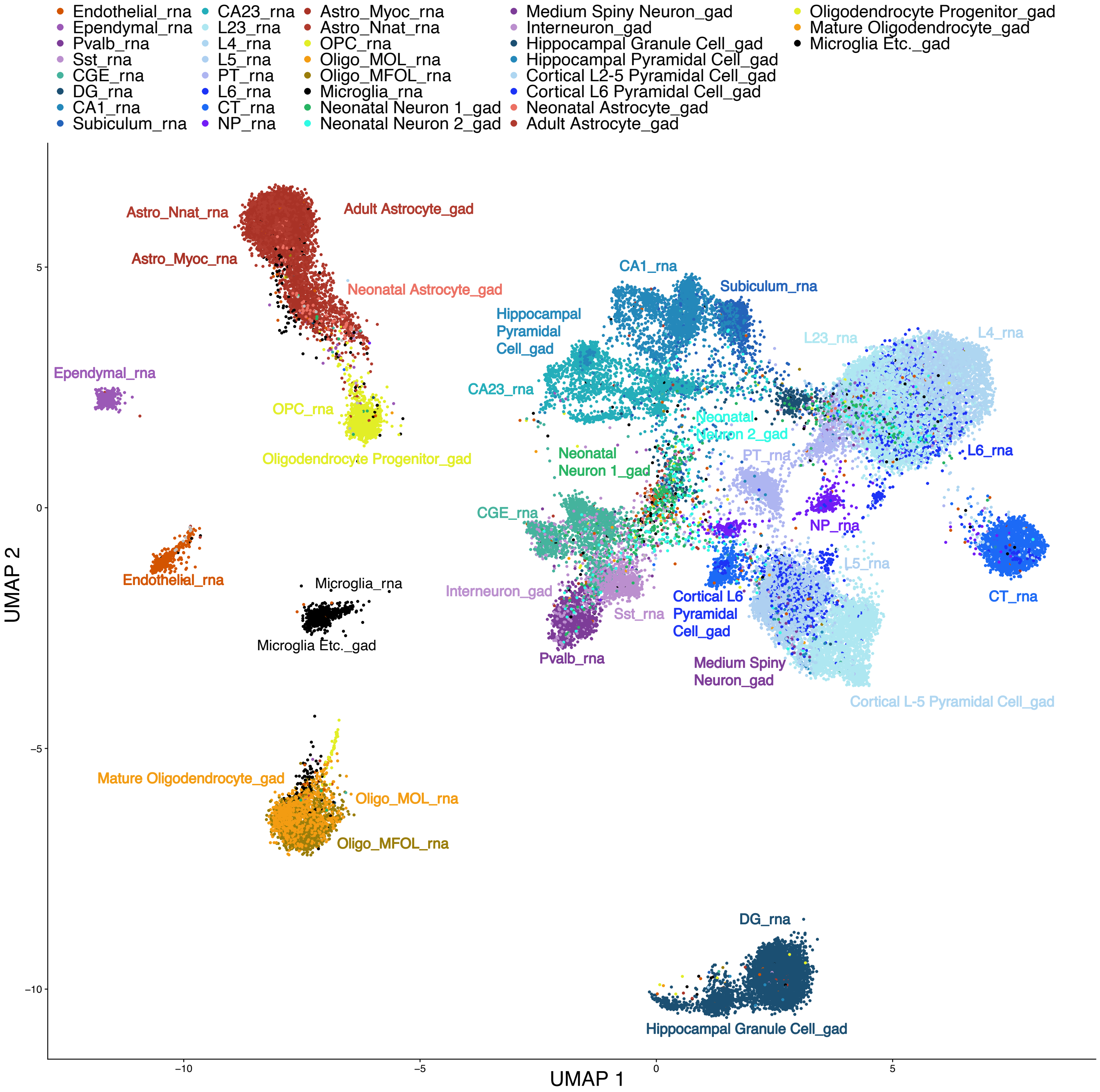


Supplementary Fig. 11: **Projection of developing mouse scHi-C data from Tan et al. 2021 onto reference low-dimensional embedding of scRNA-seq from Paired-Tag adult mouse data in Zhu et al. 2021.** Cell types from two data sets are added suffixes of ”rna” and “gad” for Paired-Tag and scHi-C data, respectively.

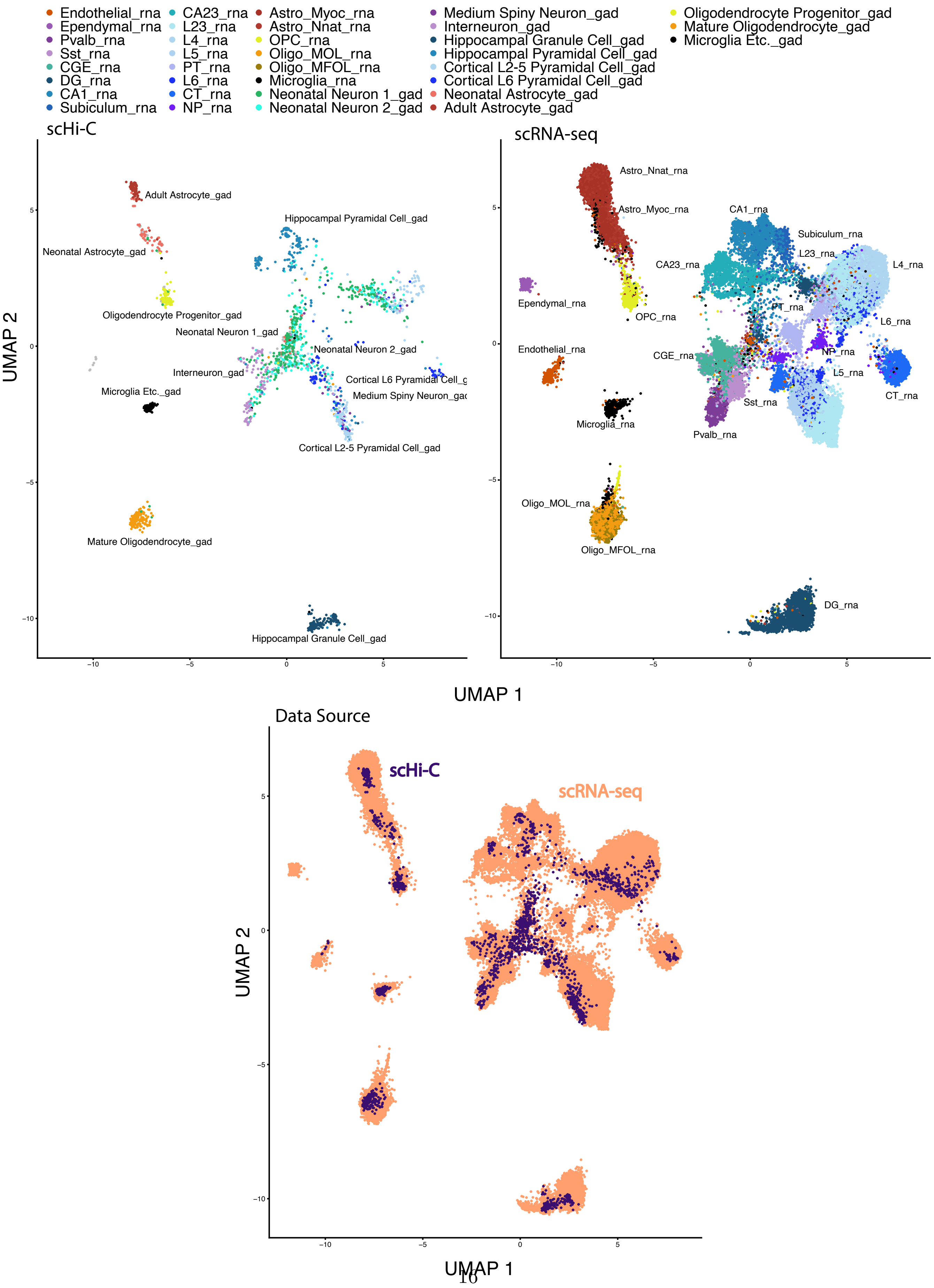




Supplementary Fig. 12: **Projection of developing mouse scHi-C data from Tan et al. 2021 onto the reference low-dimensional embedding of scRNA-seq from Paired-Tag adult mouse data in Zhu et al. 2021.** This figure expands Supplementary Figure 11 by displaying data sources on the low-dimensional embeddings.
